# Neuronal L-Type Calcium Channel Signaling to the Nucleus Requires a Novel CaMKIIα-Shank3 Interaction

**DOI:** 10.1101/551648

**Authors:** Tyler L. Perfitt, Xiaohan Wang, Jason R. Stephenson, Terunaga Nakagawa, Roger J. Colbran

**Affiliations:** Department of Molecular Physiology and Biophysics; Vanderbilt Brain Institute; Center for Structural Biology; Vanderbilt-Kennedy Center for Research on Human Development, Vanderbilt University School of Medicine, Nashville, TN, USA 37232-0615

**Keywords:** CaMKII, Cav1.3, CREB, Shank3

## Abstract

The molecular mechanisms that couple plasma membrane receptors/channels to specific intracellular responses, such as increased gene expression, are incompletely understood. The postsynaptic scaffolding protein Shank3 associates with Ca^2+^ permeable receptors or ion channels that can activate many downstream signaling proteins, including calcium/calmodulin-dependent protein kinase II (CaMKII). Here, we show that Shank3/CaMKIIα complexes can be specifically co-immunoprecipitated from mouse forebrain lysates, and that purified activated (Thr286 autophosphorylated) CaMKIIα binds directly to Shank3 between residues 829-1130. Mutation of three basic residues in Shank3 (R^949^RK^951^) to alanine disrupts CaMKII binding to Shank3 fragments *in vitro*, as well as CaMKII association with full-length Shank3 in heterologous cells. Our shRNA/rescue studies revealed that Shank3 binding to both CaMKII and L-type calcium channels (LTCCs) is required for increased phosphorylation of the nuclear CREB transcription factor induced by depolarization of cultured hippocampal neurons. Thus, this novel Shank3-CaMKII interaction is essential for the initiation of a specific long-range signal from plasma membrane LTCCs to the nucleus that is required for activity-dependent changes in neuronal gene expression during learning and memory.

## INTRODUCTION

Neuronal activation leads to depolarization and Ca^2+^ influx, stimulating multiple intracellular signaling pathways that are essential for normal brain functions. For example, Ca^2+^-dependent phosphorylation of the nuclear transcription factor CREB at Ser^133^ leads to the transcription of immediate-early genes encoding multiple proteins (e.g., *c-fos*, BDNF, homer1a) that play key roles in learning and memory (Bading, 2013, Benito et al., 2011, Dolmetsch, 2003, Flavell & Greenberg, 2008). Multiple neuropsychiatric disorders are associated with disruptions in activity-dependent gene expression (Ebert & Greenberg, 2013, Gallo et al., 2018), and these disorders have been linked to mutations in Ca^2+^ signaling proteins, such as L-type calcium channels (LTCCs) and calcium/calmodulin-dependent protein kinase II (CaMKII) (Akita et al., 2018, Chia et al., 2018, Dick et al., 2016, Kury et al., 2017, Limpitikul et al., 2016, Moon et al., 2018, Nyegaard et al., 2010, Pinggera et al., 2015, Pinggera et al., 2017, Pinggera & Striessnig, 2016, Proietti Onori et al., 2018, Stephenson et al., 2017). For example, Timothy Syndrome is caused by mutations in the Cav1.2 LTCC a1 subunit that disrupt neuronal excitation-transcription (E-T) coupling (Li et al., 2016), contributing to the neurobehavioral symptoms of this complex multi-system disorder, including autism spectrum disorder (ASD). Recent studies have shown that the initiation of LTCC-dependent E-T coupling requires the recruitment of multiple CaMKII holoenzymes to a nanodomain close to LTCCs (Ma et al., 2014, Wheeler et al., 2008). Moreover, a CaMKII mutation linked to intellectual disability disrupts neuronal E-T coupling (Cohen et al., 2018). However, the molecular mechanisms underlying the organization and function of this LTCC nanodomain remain incompletely understood.

Multiple CaMKII isoforms have critical roles in neuronal signaling and plasticity. Ca^2+^/calmodulin (CaM) binding to 12-subunit CaMKII holoenzymes stimulates intersubunit autophosphorylation at Thr286 (in CaMKIIα) in neurons, a key mechanism underlying learning and memory (reviewed in (Hell, 2014, Lisman et al., 2012, Shonesy et al., 2014)). Thr286-autophosphorylated CaMKII can remain autonomously active after the initial Ca^2+^ influx dissipates and Ca^2+^/CaM dissociates (Lai et al., 1986, Miller & Kennedy, 1986, Miller et al., 1988), leading to sustained phosphorylation of downstream targets. Activated CaMKII also interacts with a number of other synaptic proteins. For example, activated CaMKII binding to the GluN2B of the NMDA-type glutamate receptor is important in targeting CaMKII to dendritic spines and for normal synaptic plasticity (Bayer et al., 2001, Bayer et al., 2006, Halt et al., 2012, Strack & Colbran, 1998). The neuronal scaffolding protein densin can bind to both Cav1.3 LTCCs and CaMKII, acting as a scaffold for CaMKII to suppress Ca^2+^-dependent inactivation of Cav1.3, thereby facilitating Ca^2+^ entry (Jenkins et al., 2010, Jiao et al., 2011). In addition, a recent study showed that CaMKII binding to the mGlu5 metabotropic glutamate receptor modulates the mobilization of intracellular Ca^2+^ stores (Marks et al., 2018). This emerging evidence supports the hypothesis that direct interactions of CaMKII with CaMKII-Associated Proteins (CaMKAPs) are important for synaptic signaling.

Our lab recently conducted a proteomics study to identify novel CaMKAPs, revealing that Shank3 is among the most abundant proteins detected in synaptic CaMKII complexes (Baucum et al., 2015). Canonically, the full-length Shank3 protein contains an N-terminal ankyrin repeat domain, SH3 and PDZ domains, a proline-rich region with binding sites for homer and cortactin, and a C-terminal SAM domain that mediates oligomerization (Naisbitt et al., 1999). Diverse mutations in Shank3 are strongly linked to ASD and schizophrenia (Leblond et al., 2014, Soler et al., 2018), while a chromosomal deletion causes haploinsufficiency of the *SHANK3* gene in 22q13 deletion syndrome (Phelan-McDermid Syndrome), another neurodevelopmental disorder associated with ASD (Harony-Nicolas et al., 2015). Indeed, Shank3 is crucial for the formation and stabilization of excitatory synapses (Verpelli et al., 2011). Over a dozen unique Shank3 mutant mouse lines display different combinations of deficits in synaptic transmission, social behavior, and learning (Monteiro & Feng, 2017). Since interaction of the Shank3 PDZ domain with a C-terminal PDZ-binding motif in Cav1.3 LTCCs was reported to be involved in both clustering of LTCCs in neuronal dendrites and LTCC signaling to increase CREB phosphorylation (Zhang et al., 2005), we hypothesized that a direct interaction of CaMKII with Shank3 is also important in CaMKII recruitment to the LTCC nanodomain that is required for E-T coupling.

Here we identify a novel binding site for CaMKII in Shank3 between the PDZ domain and homer-binding motif and show that CaMKII autophosphorylation at Thr286 is required for this interaction. Using site-directed mutagenesis, we identified three residues in Shank3 that are critical for this interaction. Mutation of these residues in full-length Shank3 disrupts the co-immunoprecipitation of CaMKII from heterologous cell lysates, and the colocalization of CaMKII in heterologous cells. In addition, this mutation disrupts LTCC-CREB signaling in primary hippocampal neurons.

## RESULTS

### CaMKIIα and Shank3 interact in the mouse forebrain

Our previous proteomics study detected numerous peptides originating from the Shank3 protein in CaMKII immune complexes isolated from Triton-insoluble synaptic fractions of mouse forebrain (Baucum et al., 2015). To extend this observation, we first compared the distribution of Shank3 and CaMKII across cytosolic (S1), Triton-soluble membrane (S2), and Triton-insoluble synaptic (P2) fractions isolated from mouse forebrain extracts. The Shank3 antibody detected two major bands in whole mouse brain extracts - the expected ~180 kDa band, plus a ~125 kDa band. Both Shank3 bands were undetectable in S1 or S2 fractions and were relatively enriched in the P2 fraction, similar to other synaptic proteins, such as PSD-95 (Fig. 1A). Since *Shank3* undergoes complex transcriptional and post-transcriptional regulation through intragenic promoters and alternative splicing (Waga et al., 2013, Wang et al., 2011), the two bands may represent different Shank3 variants. Alternatively, the smaller protein may be a proteolytic fragment of the full length Shank3, consistent with observed variability of the ratio of these two bands between independent experiments (compare with input lanes in Figs. 1B, 1C). In contrast, similar levels of CaMKII were detected in the S2 and P2 fractions, with lower levels in S1, consistent with our prior studies (Gustin et al., 2011). Thus, a subpopulation of CaMKII and essentially all of the Shank3 are present in mouse brain subcellular fractions enriched in synaptic proteins.

**Figure 1.**
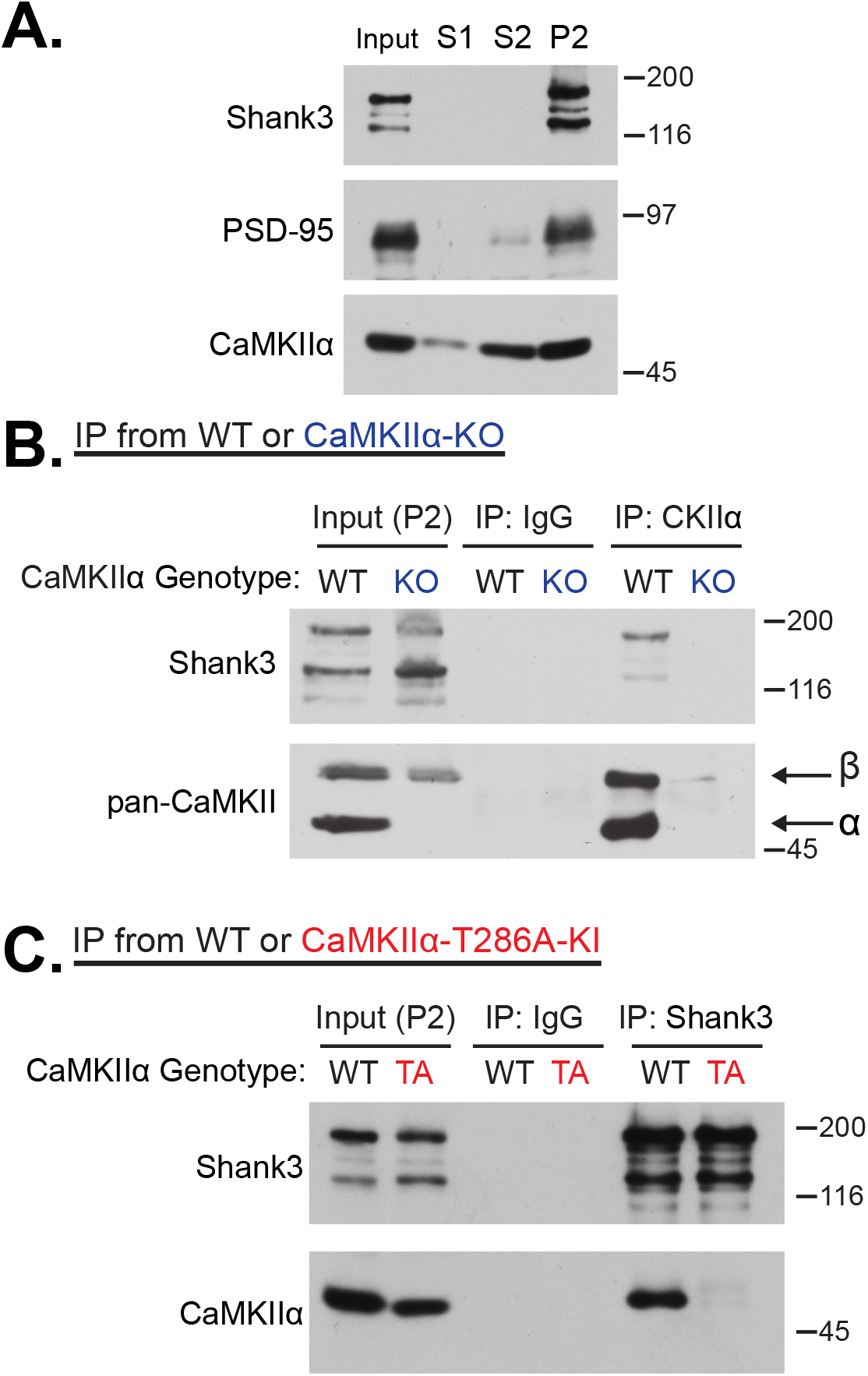
Reciprocal co-immunoprecipitation of Shank3 and CaMKIIα from mouse forebrain extracts. **A.** Cytosolic (S1), membrane-associated (S2), and synaptic (P2) subcellular fractions were immunoblotted for localization of Shank3 (Cell Signaling antibody), PSD-95, and CaMKIIα. Shank3 is primarily localized in P2 fractions, which is also enriched for CaMKIIα. **B.** Synaptic (P2) fractions from WT or CaMKIIα-KO mouse forebrains were immunoprecipitated using control IgG or CaMKIIα-specific antibodies, and immunoblotted using Shank3 (Cell Signaling antibody) and pan-CaMKII antibodies. The CaMKIIα and CaMKIIβ bands are indicated by arrows. Shank3 was not detected in samples isolated from CaMKIIα-KO mice. **C.** Synaptic (P2) fractions from WT or CaMKIIα^T286A^ mouse forebrains were immunoprecipitated using control IgG or Shank3 (Bethyl) antibodies, and immunoblotted using Shank3 (Cell Signaling) and CaMKIIα antibodies. Co-precipitated CaMKIIα is significantly reduced in CaMKIIα^T286A^ mice (93±4% reduction compared to WT, n=3, p=0.0008, one sample Student’s t-test with equal variance compared to theoretical value of 100). All immunoblots are representative of 3 biological replicates.

In order to investigate the specificity of Shank3 association with CaMKII, we incubated synaptic P2 fractions, isolated in parallel from wild-type or CaMKIIα knockout (CaMKIIα-KO) littermates, with a control IgG antibody or a monoclonal CaMKIIα-specific antibody. Immunoblotting revealed that similar levels of Shank3 and CaMKIIβ were present in WT and CaMKIIα-KO P2 fractions. Shank3 was present in CaMKIIα immune complexes isolated from WT mouse forebrain, but not from CaMKIIα-KO mouse forebrain, and not in any samples isolated using a control IgG (Fig. 1B). Interestingly, the ratio of the higher and lower molecular weight Shank3 bands was consistently higher in the CaMKII immune complexes than in the P2 input, perhaps suggesting that CaMKII preferentially interacts with the larger Shank3 protein. These data confirm that Shank3 specifically associates with CaMKII complexes in mouse brain extracts.

We next tested for reciprocal co-immunoprecipitation of CaMKII with complexes isolated using a Shank3 antibody. Since interactions between CaMKII and other proteins are often enhanced by Thr286 autophosphorylation, we compared the association of CaMKIIα with Shank3 in synaptic P2 fractions isolated from WT mice and homozygous CaMKIIα^T286A^ mice with a knock-in mutation of Thr286 to Ala, preventing CaMKII regulation by Thr286 autophosphorylation (Giese et al., 1998). While the genotype did not appear to affect the levels of Shank3 in the inputs or in the immune complexes, as detected by immunoblotting using a different Shank3 antibody, there were significantly lower levels of CaMKIIα in Shank3 immune complexes isolated from homozygous CaMKIIα^T286A^ mice than from WT littermates (Fig. 1C) (93±4% reduced compared to WT, p=0.0008, one sample Student’s t-test with equal variance compared to theoretical value of 100, *n* = 3). In combination, these data indicate that Shank3 interacts directly or indirectly with CaMKII in a Triton-insoluble synaptic fraction of mouse brain, and that this association is sensitive to the levels of Thr286 autophosphorylation of CaMKII.

### Activated CaMKIIα directly binds to Shank3 (829-1130)

To determine whether CaMKIIα directly interacts with Shank3, we first expressed and purified a series of six non-overlapping GST fusion proteins spanning the full length of Shank3 (Figs. 2A, 2B). Some of the purified proteins contained proteolytic degradation fragments and the full-length proteins are denoted by asterisks in Fig. 2B. Each protein was incubated with purified Thr286-autophosphorylated CaMKIIα and complexes were isolated using glutathione agarose. A GST fusion protein containing the CaMKII-binding domain of the NMDA receptor GluN2B subunit (residues 1260-1309), a well-established CaMKAP (Strack & Colbran, 1998), was used as a positive control. Similar amounts of activated CaMKIIα bound to a GST-Shank3 fusion protein containing residues 829-1130 (GST-Shank3 #4 in the figure) and to GST-GluN2B, but there was no consistently detectable interaction with any other Shank3 fragment (Fig. 2B). Notably, CaMKII activation (Thr286 phosphorylation) is essential for binding to GST-Shank3 (829-1130) (Fig. 2C). Inactive CaMKIIα also did not bind to any of the other GST-Shank3 proteins compared to GST negative control (Figure EV1). In combination, these data show that activated CaMKII can directly interact with a central domain in the Shank3 protein.

**Figure 2.**
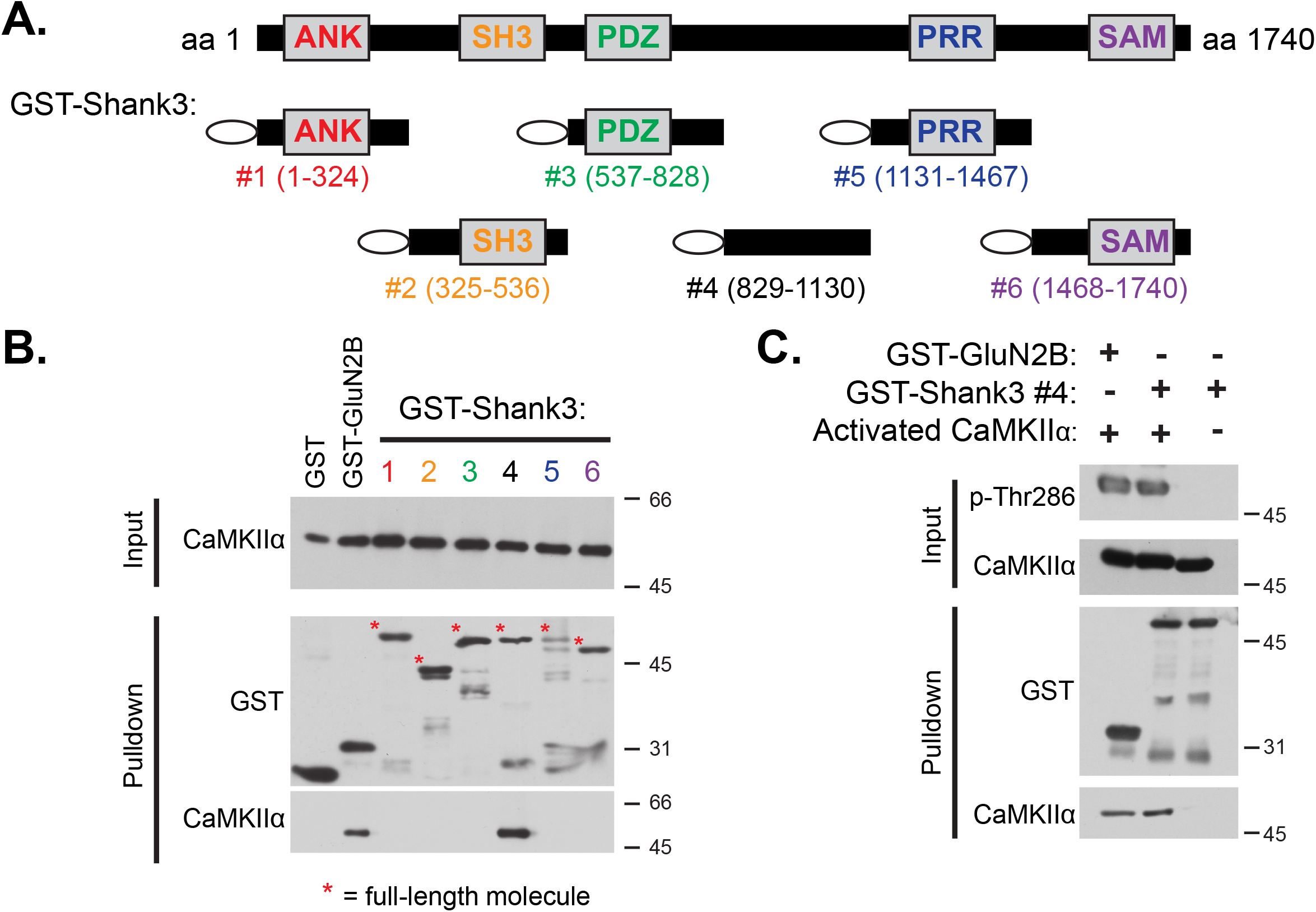
Activated CaMKIIα specifically binds to Shank3 (829-1130). **A.** Domain structure of full-length Shank3 and the six GST-Shank3 fusion proteins used in these studies that span the entire Shank3 protein. Canonical Shank3 domains are depicted as gray boxes and residue numbers are listed in parentheses. ANK = ankyrin-rich repeats, SH3 = Src homology 3 domain, PDZ = PSD95/Dlg1/zo-1 domain, PRR = proline-rich region containing binding sites for Homer and Cortactin, SAM = Sterile alpha motif involved in multimerization of Shank3. **B.** Glutathione agarose co-sedimentation assay shows that pre-activated (Thr286-autophosphorylated) CaMKIIα specifically binds to GST-Shank3 #4 (829-1130) and positive control GST-GluN2B (1260-1309). Full-length GST fusion proteins denoted with asterisks on the GST immunoblot. **C.** Glutathione agarose co-sedimentation assay shows that inactive CaMKIIα does not bind to GST-Shank3 #4 (829-1130); *in vitro* binding required pre-activation of CaMKIIα. Immunoblots are representative of 3-4 biological replicates.

### A Shank3 tri-basic residue motif is required for CaMKII binding

Residues 829-1130 of Shank3 are between the canonical PDZ domain and the proline rich motif, and the functional role(s) of this region is poorly understood (Naisbitt et al., 1999, Tu et al., 1999). In order to identify CaMKII-binding determinants within this domain, we initially generated and purified three smaller GST-Shank3 fusion proteins (Fig. 3A). Similar amount of activated CaMKII bound to GST-Shank3 (931-1014) (protein 4b in the figure) and GST-Shank3 (829-1130), but there was no detectable interaction with GST-Shank3 proteins containing residues 829-930 (protein 4a) or residues 1015-1130 (protein 4c) (Fig. 3B). Examination of the amino acid sequence of Shank3 residues 931-1014 revealed a region sharing similarity with recently-characterized CaMKII binding domains in the N-terminal domain of the LTCC Cav1.3 a subunit (Wang et al., 2017) and the C-terminal domain of mGlu5 (Marks et al., 2018) (Fig. 3A). Mutation of tri-basic residue motifs in Cav1.3 (^83^Arg-Lys-Arg^85^) and mGlu5 (^866^Lys-Arg-Arg^868^) prevented CaMKIIα binding. Therefore, we mutated the conserved Shank3 ^949^Arg-Arg-Lys^951^ motif to Ala residues within GST-Shank3 (829-1130); this RRK/AAA mutation essentially abrogates the binding of activated CaMKIIα (Fig. 3C). These data indicate that ^949^Arg-Arg-Lys^951^ in Shank3 are essential for the direct binding of activated CaMKII *in vitro*.

**Figure 3.**
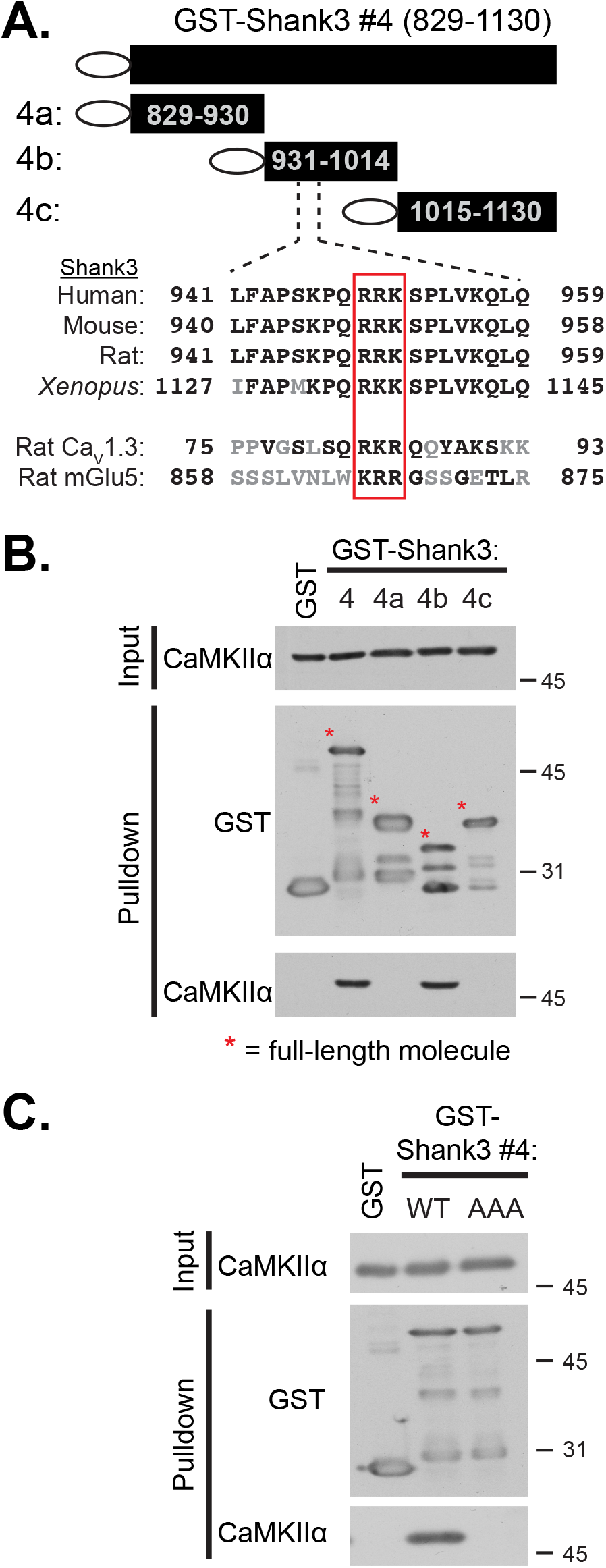
Characterization of the CaMKII-binding motif in Shank3. **A.** *Top*, Diagram of 3 truncations used to map the CaMKII interaction site within GST-Shank3 #4 (829-1130). *Bottom*, Sequence alignment of human Shank3 residues 941-959 with the corresponding Shank3 residues in other species and the CaMKII-binding domain in the N-terminal domain of the *Rattus norvegicus* Cav1.3 a1 subunit (Wang et al., 2017) and the C-terminal tail of the *Rattus norvegicus* mGlu5 (Marks et al., 2018), with conserved residues in black and dissimilar residues in grey. The conserved tri-basic residue motif is highlighted in red box. **B.** Glutathione agarose co-sedimentation assay comparing binding of activated CaMKIIα to GST-Shank3 #4 (829-1130) and 3 nonoverlapping fragments (4a, 4b, and 4c). Full-length GST fusion proteins denoted with asterisks. GST-Shank3 #4 (829-1130) and #4b (931-1014) bind similar amounts of activated CaMKIIα, but there is no detectable binding to the other Shank3 fragments. **C.** Mutation of amino acids ^949^RRK^951^ to AAA in GST-Shank3 #4 (829-1130) blocks CaMKIIα binding in glutathione agarose co-sedimentation assay (98±4% reduced compared to WT, n=3, p=0.0007, one-sample Student’s t-test with equal variance compared to theoretical value of 100). All immunoblots are representative of 3 biological replicates.

### The RRK/AAA mutation specifically disrupts binding of CaMKII to full-length Shank3

In order to determine whether the ^949^Arg-Arg-Lys^951^ motif is essential for association of CaMKII with full-length Shank3, we generated the RRK/AAA mutant in GFP-tagged full-length Shank3 (GFP-Shank3-AAA). WT CaMKIIα was co-expressed with GFP, GFP-Shank3-WT, or GFP-Shank3-AAA in HEK293T cells and a GFP antibody was used for immunoprecipitation after the addition of excess Ca^2+^/CaM to activate CaMKIIα in the cell lysates. Similar robust GFP protein signals were detected in samples immunoprecipitated from each lysate, but CaMKIIα was only detected in immune complexes containing GFP-Shank3-WT, but not GFP control or GFP-Shank3-AAA complexes (Fig. 4A).

**Figure 4.**
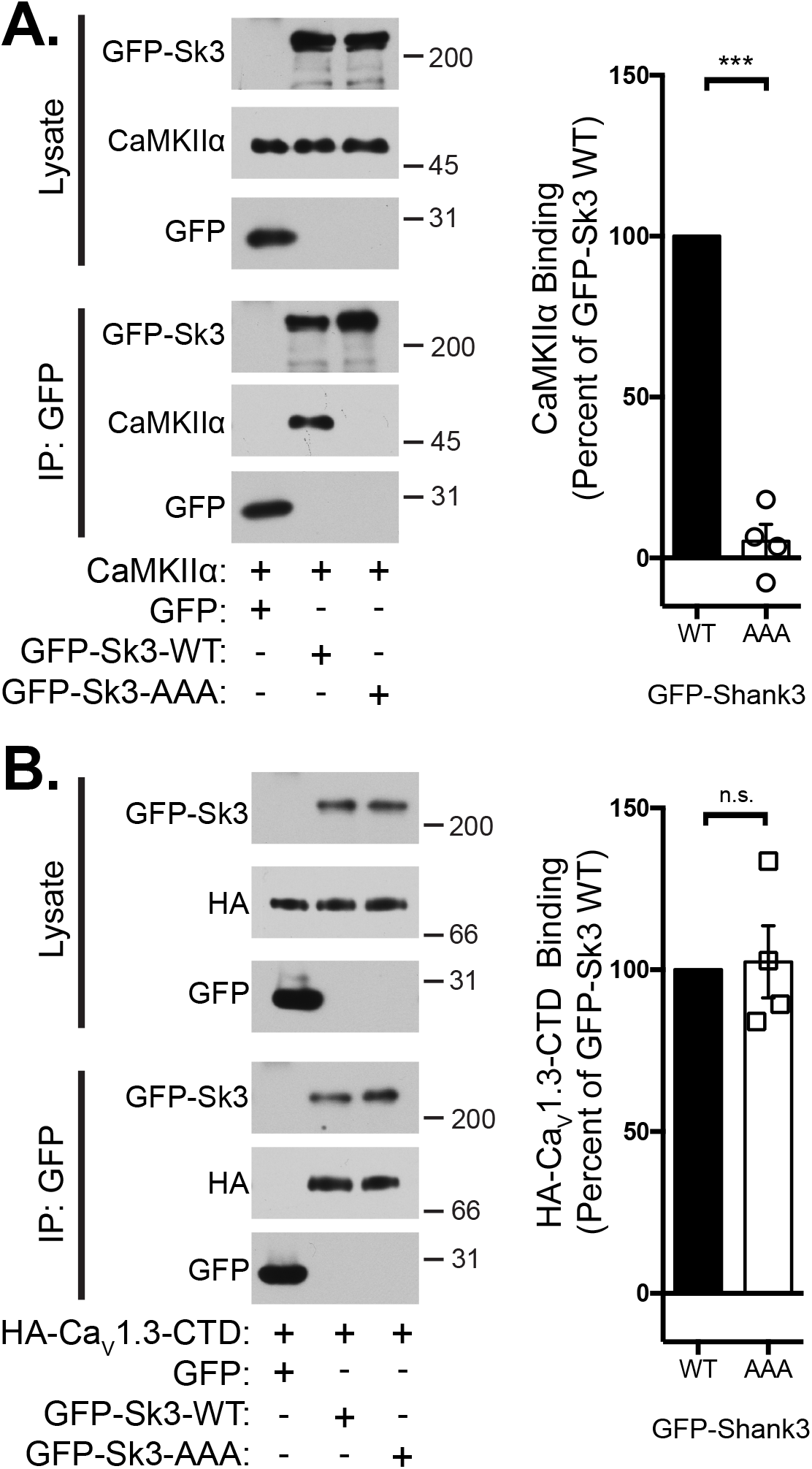
The ^949^RRK^951^ to AAA mutation in Shank3 disrupts the association with CaMKIIα, but does not with the Cav1.3 C-terminal domain. **A.** Soluble fractions of HEK293T cells expressing CaMKIIα with GFP or GFP-Shank3 (WT or ^949^RRK^951^ to AAA mutant) were immunoprecipitated using a GFP antibody. Co-precipitation of CaMKIIα with GFP-Shank3 AAA is significantly reduced by 95±10% (n=4) compared to GFP-Shank3 WT (*** p=0.0004, one-sample unpaired Student’s t-test with equal variance compared to a theoretical value of 100). **B.** Soluble fractions of HEK293T cells expressing GFP or GFP-Shank3 (WT or AAA) with the HA-tagged C-terminal domain of the Cav1.3 a1 subunit (HA-Cav1.3-CTD), were immunoprecipitated as in panel A. The AAA mutation has no significant effect on the co-precipitation of HA-Cav1.3-CTD (n=4; p=0.84). All immunoblots are representative of 4 biological replicates. Bar graphs show the mean ± SEM.

To test whether the RRK/AAA mutation has a specific effect on CaMKII binding to Shank3, we performed a similar co-immunoprecipitation experiment using an HA-tagged C-terminal domain (CTD) of the L-type calcium channel Cav1.3 α subunit, which has been previously shown to interact with the nearby PDZ domain (residues 572-661) of Shank3 (Zhang et al., 2005). Lysates of HEK293T cells expressing the HA-Cav1.3-CTD and either GFP, GFP-Shank3-WT, or GFP-Shank3-AAA were immunoprecipitated using a GFP antibody. Similar amounts of the HA-Cav1.3-CTD co-immunoprecipitated with both GFP-Shank3-WT and GFP-Shank3-AAA immune complexes, but not with GFP alone (Fig. 4B). Thus, interactions with the Shank3 PDZ domain are not affected by mutation of the ^949^Arg-Arg-Lys^951^ motif, indicating that this mutation does not have broader (non-specific) effect on the interactions of other proteins with GFP-Shank3.

### RRK/AAA disrupts colocalization between GFP-Shank3 and mApple-CaMKIIα in striatal progenitor cells

Shank3 and CaMKIIα are expressed at high levels in neurons. Both proteins are strongly localized to dendritic spines (Boeckers et al., 2005, Shen & Meyer, 1999), but it is well established that other CaMKAPs have important roles in bulk targeting of CaMKII to spines (such as the NMDA receptor GluN2B subunit (Bayer et al., 2006, Halt et al., 2012)). To avoid potentially confounding effects of GluN2B or other CaMKAPs on the interpretation of data from studies examining the association (colocalization) of Shank3 and CaMKII in neurons, we investigated the impact of Shank3 on CaMKII localization in striatal ST*Hdh*^+/+^ progenitor cells, which extend neurite-like outgrowths in response to a pharmacological stimulation that induces a neuron-like differentiation program (see Methods) (Trettel et al., 2000). We first transfected cells with GFP-Shank3-WT and soluble mApple (mAp). After differentiation, GFP-Shank3-WT formed large puncta along the cell perimeter and was localized to the neurite-like outgrowths, while mAp was diffusely localized throughout the cell, with no clearly discernable mAp punctae that overlapped with the GFP punctae (Fig. 5A, left; green arrowheads). Co-expression of mAp-CaMKIIα with GFP-Shank3-WT resulted in an uneven distribution of the mAp signal within both the cell body and neurite-like outgrowths, with mAp punctae consistently, if only partially, overlapping with GFP punctae (Fig. 5A, center, white arrowheads). However, when co-expressed with GFP-Shank3-AAA, the mAp-CaMKIIα signal appeared to be unevenly concentrated in the cell body with no obvious overlap with the GFP signal, and little mAp fluorescence detected in the processes (Fig. 5A, right, green arrowheads). We quantified the colocalization of the mAp and GFP signals across multiple cells co-transfected with these combinations of fluorescent proteins using image correlation analysis (Li et al., 2004) (Fig. 5B). There was minimal colocalization of soluble mAp with GFP-Shank3-WT (ICQ: 0.07±0.01), whereas mAp-CaMKIIα was strongly colocalized with GFP-Shank3-WT (ICQ: 0.31±0.02). In contrast, mAp-CaMKIIα and GFP-Shank3-AAA displayed minimal colocalization (ICQ: 0.09±0.02). The ICQ for mAp-CaMKIIα and GFP-Shank3-WT is significantly greater than that of either mAp and GFP-Shank3-WT or of mAp-CaMKIIα and GFP-Shank3-AAA (Fig. 5B) (1-way ANOVA, F_(2,70)_ = 39.55, p < 0.0001. Tukey’s post-hoc test, **** p < 0.0001). Thus, these data indicate that the RRK/AAA mutation in Shank3 not only disrupts binding with CaMKII *in vitro*, but also the association of CaMKII with Shank3 in intact cells.

**Figure 5.**
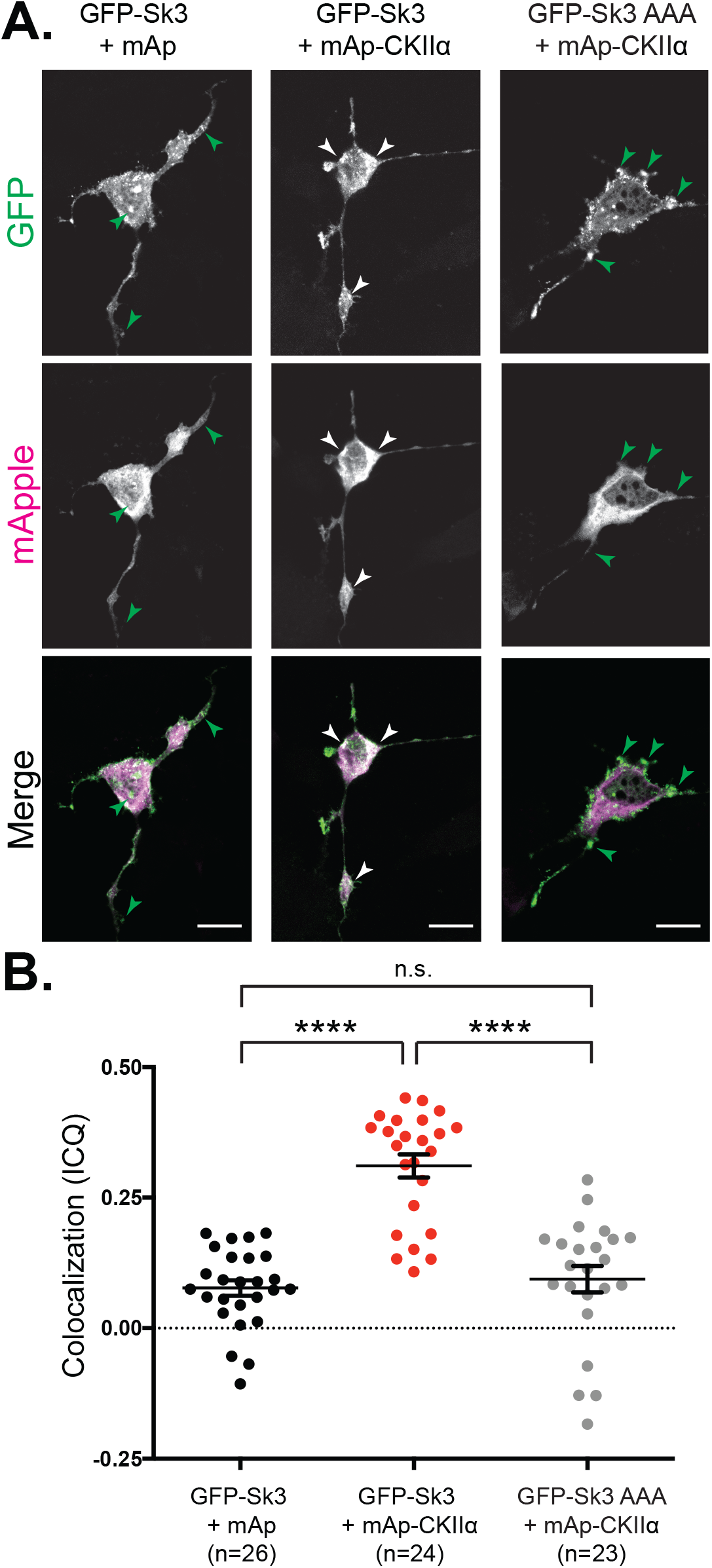
The ^949^RRK^951^ to AAA mutation in Shank3 disrupts colocalization with CaMKIIα in striatal progenitor cells. **A.** Representative images of differentiated striatal ST*Hdh*^+/+^ progenitor cells expressing GFP-Shank3 (WT or ^949^RRK^951^ to AAA mutant) with mApple (mAp) or mAp-CaMKIIα. GFP punctae without corresponding mAp signal are denoted by green arrowheads, while GFP punctae overlapping mAp signal are denoted by white arrowheads. *Scale bars* 10 μm. **B.** ICQ (Intensity Correlation Quotient) analysis comparing the distribution of GFP and mAp signals in striatal cells. ICQ values for GFP-Shank3 and mAp-CaMKIIα (mean±SEM: 0.31±0.02) are significantly higher than GFP-Shank3 and mApple (0.07±0.01) or GFP-Shank3 AAA and mAp-CaMKIIα (0.09±0.02) (1way ANOVA, F(2,70) = 39.55, p < 0.0001. Tukey’s post-hoc test, **** p < 0.0001). Each data point plotted indicates the ICQ value from a single cell, with 7-12 cells analyzed from each of 3 independent cultures/transfections.

### Shank3 is required for LTCC- and CaMKII-mediated signaling to the nucleus

To begin probing the functional role of Shank3 binding to CaMKII, we used a previously established paradigm to induce LTCC- and CaMKII-dependent Ser^133^ phosphorylation of the nuclear transcription factor CREB in primary hippocampal neurons (Li et al., 2016, Wang et al., 2017, Wheeler et al., 2008). Neurons were pre-incubated in 5 mM K^+^ Tyrode’s solution (5K) containing APV and CNQX to block the activation of NMDA- and AMPA-type glutamate receptors and with tetrodotoxin (TTX) to inhibit voltage-dependent sodium channels (Fig. 6A). Neuronal depolarization, induced by replacing the solution with 40 mM K^+^ Tyrode’s solution (40K) in the presence of APV, CNQX, and TTX for 90 seconds, resulted in a significant increase in nuclear staining of a phospho-Ser^133^-specific CREB antibody (pCREB intensity) (Fig. 6B, black bars). It is well established that this depolarization-induced increase in pCREB staining can be completely disrupted by the selective LTCC blocker nimodipine (10 μM) (Wheeler et al., 2012). Moreover, this depolarization protocol results in significant LTCC-dependent expression of the c-fos protein 3hr after washing off the 40K stimulation solution (Fig. EV2).

**Figure 6.**
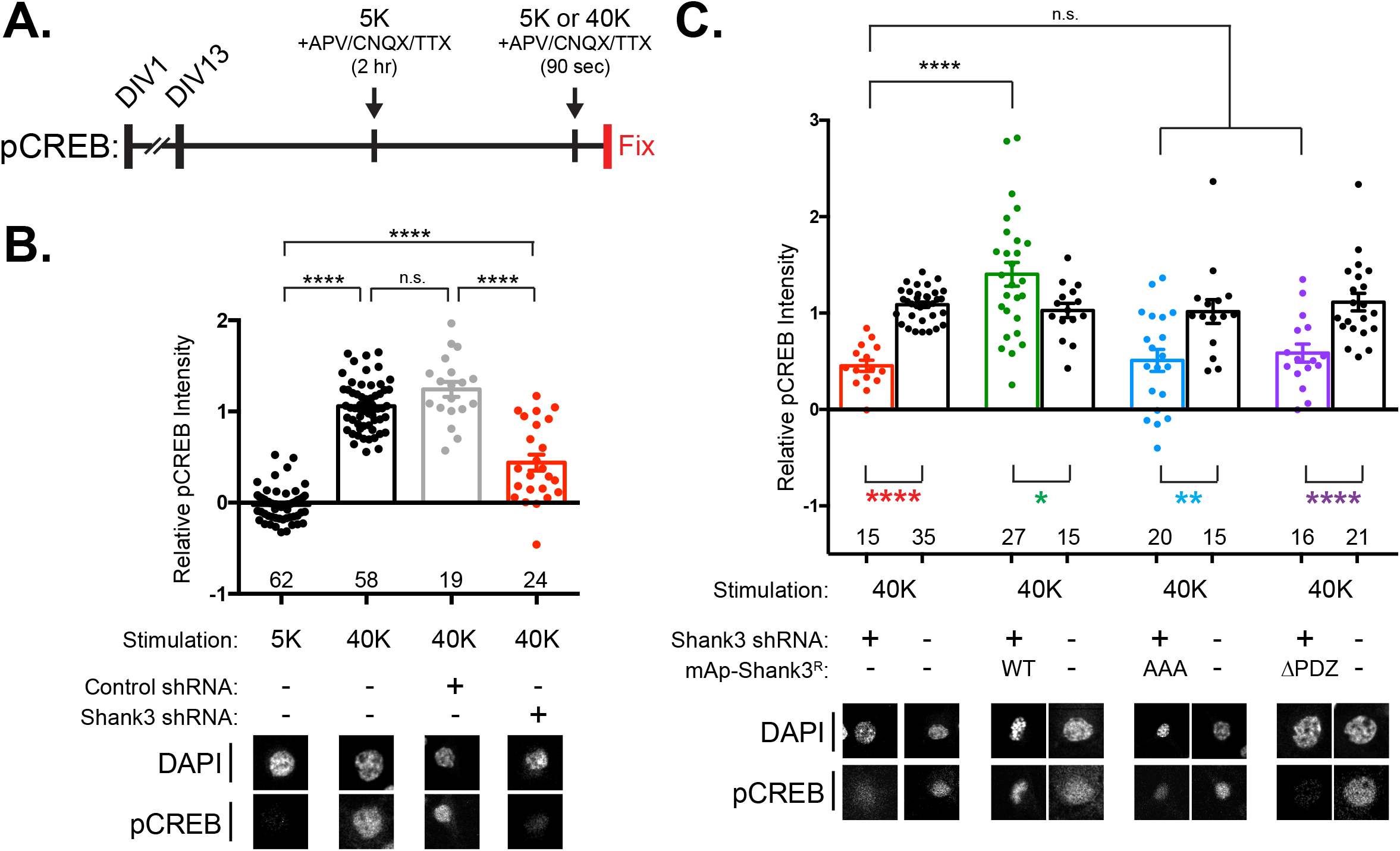
The ^949^RRK^951^ to AAA mutation in Shank3 disrupts LTCC signaling to CREB in neurons. **A.** Schematic of experimental protocol. Primary hippocampal neurons (DIV13) were incubated with inhibitors of NMDA- and AMPA-type glutamate receptors and voltage-gated sodium channels (APV, CNQX, and TTX) in 5 mM KCl (5K) Tyrode’s solution for 2 hours, and then treated with 5K (control) or 40 mM KCl (40K: depolarizing) Tyrode’s solution for 90 seconds before fixation. Neurons were then stained using antibodies to pSer133-CREB and DAPI. **B.** The 40K stimulation induced a robust increase in nuclear CREB phosphorylation, which is significantly reduced in cells expressing the Shank3 shRNA (red bar), but not control shRNA (grey bar) (1-way ANOVA, F(3,159) = 189.7, p < 0.0001. Tukey’s post-hoc test, **** p < 0.0001). **C.** The reduction of nuclear CREB phosphorylation in an independent set of Shank3 shRNA neurons was rescued by co-expressing shRNA-resistant mAp-Shank3^R^-WT (green bar), but not mAp-Shank3^R^-AAA (blue bar) or mAp-Shank3^R^-ΔPDZ (purple bar) (2-way ANOVA with two factors (Transfection, Rescue), Transfection F(1,156) = 22.70, Rescue F(1,156) = 10.14, Interaction F(1,156) = 12.29, all p < 0.001. Tukey’s post-hoc test, **** p < 0.0001). In each transfection, nearby non-transfected cells were analyzed and no effect on CREB phosphorylation was observed (black bars). Each data point represents analysis of single cells accumulated from 3-5 independent neuronal cultures/transfections. All confocal images show a 40 x 40 μm area. The bar graph reports the mean ± SEM.

We first tested whether Shank3 expression is required for this specific signaling mechanism using an shRNA approach. A control shRNA, which does not target any gene transcript (Boudkkazi et al., 2014), had no effect on pCREB intensity in stimulated neurons (Fig. 6B, gray bar). However, transfection with an shRNA targeting Shank3 (Verpelli et al., 2011)(Fig. EV3A) significantly reduced the depolarization-stimulated pCREB staining to 44±8% of the staining intensity in control neurons (Fig 6B, red bar) (1-way ANOVA, F_(3,159)_ = 189.7, p < 0.0001. Tukey’s post-hoc test, **** p < 0.0001). Thus, LTCC-dependent increases in CREB phosphorylation are at least partially dependent on the presence of Shank3.

We then investigated whether co-transfection with shRNA-resistant Shank3 constructs (mAp-Shank3^R^) (Fig. EV3A), could rescue the effect of shRNA knockdown on CREB phosphorylation following stimulation with 40 mM KCl. The decrease in pCREB staining in neurons transfected with Shank3 shRNA alone (Fig. 6C, red bar, 45±6% of control in this independent set of 3 experiments), was rescued by co-expression of mAp-Shank3^R^-WT (Fig. 6C, green bar, 140±12% of control), but not by co-expression of mAp-Shank3^R^ with the AAA mutation (mAp-Shank3^R^-AAA) (Fig. 6C, blue bar, 51±11% of control). Similarly, the co-expression of mAp-Shank3^R^ with an internal deletion of the PDZ domain (mAp-Shank3^R^-ΔPDZ), which is unable to bind to the C-terminal domain of the Cav1.3 LTCC (Zhang et al., 2005) (Fig. EV3B), also failed to rescue pCREB staining (Fig. 6C, purple bar, 59±9% of control). In all of these studies, we confirmed that nearby non-transfected neurons within the same culture dishes exhibited normal increases in pCREB staining intensity relative to control dishes, demonstrating that the depolarization treatment was effective in each dish of neurons (Fig. 6C, black bars) (2-way ANOVA with two factors (Transfection, Rescue), Transfection F_(1,156)_ = 22.70, Rescue F_(1,156)_ = 10.14, Interaction F_(1,156)_ = 12.29, all p < 0.001. Tukey’s post-hoc test, **** p < 0.0001). Taken together, these studies highlight the importance of Shank3 in neuronal LTCC-dependent signaling to CREB by confirming the importance of the Shank3 PDZ domain, presumably for interaction with Cav1.3 (Zhang et al., 2005), and by demonstrating that CaMKII binding to Shank3 plays a key role in this pathway.

## DISCUSSION

Our previous proteomics study indicated that the scaffolding protein Shank3 is enriched in CaMKII complexes isolated from synaptic fractions of mouse forebrain (Baucum et al., 2015). Here, we confirmed this result by immunoblotting and showed that co-immunoprecipitation is specific using forebrains from CaMKIIα-KO mice. Moreover, the amount of co-immunoprecipitation is reduced from brain lysates of CaMKIIα^T286A^ mice, which are unable to undergo CaMKIIα autophosphorylation. Direct binding of purified CaMKIIα to Shank3 residues 829-1130 also requires Thr286 phosphorylation. We showed that mutation of a tri-basic motif in Shank3 (^949^Arg-Arg-Lys^951^) to three Ala residues (AAA) completely disrupted CaMKIIα binding, but does not disrupt binding to another known Shank3 binding partner, the Cav1.3-CTD. The AAA mutation also disrupts the colocalization of mAp-CaMKIIα with GFP-Shank3 in intact striatal progenitor cells. Finally, we showed that the CaMKII-Shank3 interaction is required for LTCC- and CaMKII-dependent signaling to increase CREB phosphorylation in neuronal nuclei.

### Tri-basic residues as a new subgroup of CaMKII-binding domains

CaMKII has a wide variety of physiological functions, mediated in part by its interactions with CaMKAPs. Most CaMKAPs preferentially interact with activated CaMKII conformations, elicited by Ca^2+^/calmodulin-binding and/or Thr286 autophosphorylation. The CaMKII-binding domains of activation-dependent CaMKAPs can be classified into distinct subgroups based on sequence homology. The NMDA receptor GluN2B subunit and LTCC β1/2 subunits contain CaMKII-binding domains that share sequence homology with the auto-regulatory domain of CaMKII, and include CaMKII phosphorylation sites homologous to the Thr286 autophosphorylation site in CaMKIIα itself (Grueter et al., 2008, Strack et al., 2000). In contrast, an internal CaMKII-binding domain in densin, another synaptic scaffolding protein, shares no sequence similarity to these proteins, but resembles the naturally-occurring CaMKII inhibitor protein CaMKIIN (Jiao et al., 2011). Recent studies have identified CaMKII-binding domains containing critical tri-basic residue motifs in the N-terminal domains of Cav1.2 and Cav1.3 a1 subunits, as well as the C-terminal domain of mGlu5 (Marks et al., 2018, Wang et al., 2017). Notably, the amino acid sequence surrounding the critical CaMKII-binding tri-basic residue motif in Shank3 shares only partial homology with the sequences surrounding the CaMKII-binding tri-basic residue motifs in Cav1.3 or mGlu5 (Fig. 3A). The fact that the tri-basic residue motif alone is insufficient for CaMKII binding is demonstrated by the current observations that two GST-Shank3 proteins (#3 and #4a, respectively) contain tri-basic sequences (^707^Arg-Arg-Lys^709^ and ^919^Lys-Arg-Arg^921^, respectively), but do not bind active CaMKIIα (Figs. 2B, 3B). Further studies will be required to understand the contributions of residues surrounding the tri-basic binding motifs, as well as the potential importance of secondary or tertiary structure, to the specific binding of activated CaMKII.

### CaMKII targeting to Shank3 by CaMKIIα-Thr286 phosphorylation

Autophosphorylation of CaMKIIα at Thr286 is important for the full and persistent activation of CaMKIIα. Mice with a knock-in Ala mutation at Thr286 (CaMKIIα^T286A^) display abnormal synaptic plasticity and diverse behavioral deficits (Giese et al., 1998, Gustin et al., 2011). We found that almost no CaMKIIα-T286A co-immunoprecipitated with Shank3 from CaMKIIα^T286A^ mice (Fig. 1C). Using purified proteins to avoid potential effects of indirect interactions, we show that Shank3 is unable to bind CaMKIIα under basal, non-phosphorylated conditions (Fig. 2C). Therefore, like many other interactions, Thr286 autophosphorylation acts as a switch for CaMKII binding to Shank3 (Marks et al., 2018, Wang et al., 2017).

While it is very well established that both activated CaMKII and Shank3 are highly enriched in dendritic spines, several other synaptic/spine proteins also can bind to activated/autophosphorylated CaMKIIα. For instance, the bulk translocation of activated CaMKIIα to dendritic spines has been shown to require interactions with the C-terminal domain of the NMDAR GluN2B subunit (Bayer et al., 2006). Notably, knock-in point mutations in GluN2B that disrupt activity-dependent CaMKII binding in mice significantly reduce CaMKII targeting to PSDs, and impair both synaptic plasticity and memory recall in the Morris Water Maze (Halt et al., 2012). Therefore, we posit that Shank3 may target only a distinct subpopulation of the total postsynaptic CaMKII holoenzymes to specific complexes, consistent with the known roles of Shank3 as a modular synaptic scaffolding protein (Wang et al., 2019).

### Role of CaMKII-Shank3-Cav1.3 complex in LTCC-CREB signaling

Neuronal depolarization activates multiple signaling pathways to elicit diverse cellular responses. These include the simulation of gene transcription, which is important for long-term, activity-dependent changes in synaptic properties and behavior. In particular, diverse neuronal stimulation paradigms increase phosphorylation of the nuclear CREB transcription factor at Ser^133^ to stimulate the transcription of several immediate early genes. Importantly, knockout of CREB or repression of CREB through a CREB S133A transgene results in disruptions of long-term spatial memory and synaptic plasticity in mice (Bourtchuladze et al., 1994, Kida et al., 2002). LTCCs play an important role in stimulating CREB Ser^133^ phosphorylation and immediate early gene expression in cultured neurons and *in vivo* (Bading, 2013, Dolmetsch, 2003, Flavell & Greenberg, 2008). Thus, much attention has focused on elucidating signaling mechanisms that couple LTCC-dependent Ca^2+^ influx at the plasma membrane to the phosphorylation of CREB at Ser^133^.

LTCC Cav1.2 α1 subunits are expressed at higher levels than Cav1.3 α1 subunits in most neurons and appear to play the major role in E-T coupling in many brain regions (Striessnig et al., 2006). However, Cav1.3 LTCCs can play a major role in E-T coupling in some brain regions (Hetzenauer et al., 2006) and/or in response to weaker depolarization (Zhang et al., 2006). Interestingly, recent studies established that initiation of LTCC signaling to CREB requires a voltage-dependent conformation change in the LTCC and the creation of a local nanodomain of increased Ca^2+^ concentrations (Li et al., 2016). However, the molecular mechanisms supporting the creation of these LTCC nanodomains and their coupling to downstream signaling pathways are incompletely understood.

The C-terminal domains (CTDs) of Cav1.2 and Cav1.3 LTCCs contain canonical binding motifs for class 1 PDZ domains that exhibit distinct PDZ-binding specificity. Mutations of these PDZ-binding motifs disrupt the trafficking and clustering of both LTCC sub-types as well as LTCC signaling to increase CREB phosphorylation (Weick et al., 2003, Zhang et al., 2005). These CTD mutations might disrupt CREB signaling by interfering with the formation of the local Ca^2+^ nanodomain and/or with the conformational changes in the LTCC (Li et al., 2016). The CTD of Cav1.3, but not Cav1.2, was shown to interact with the PDZ domains of Shank1 or Shank3 (Zhang et al., 2005), but the specific roles of Shank in LTCC-dependent E-T coupling have not been investigated. Thus, our data significantly extend these prior findings by showing that maximal LTCC coupling to CREB phosphorylation requires Shank3 expression, and that deletion of the PDZ domain prevents the rescue of LTCC-CREB coupling by shRNA-resistant Shank3. These data are consistent with the hypothesis that Cav1.3 interaction with the Shank3 PDZ domain is important for creation of a Cav1.3 nanodomain that is required for initiation of E-T coupling. The residual E-T coupling following Shank3 knockdown that we observed may be mediated by Shank3-independent Cav1.2 LTCC signaling, although our data cannot preclude contributions from low levels of residual Shank3 expression. Nevertheless, these observations provide further support for the importance of Shank3 for LTCC signaling to the nucleus, at least under these conditions.

Initiation of LTCC-CREB signaling also requires CaMKII recruitment to the LTCC nanodomain (Wheeler et al., 2008). Our recent studies uncovered a direct interaction between the L-type Ca^2+^ channel (LTCC) Cav1.3 α1 subunit N-terminal domain and CaMKII that is necessary for LTCC Ca^2+^ influx to trigger phosphorylation of the transcription factor CREB at Ser^133^ (Wang et al., 2017). However, the present results show that LTCC-CREB signaling also requires CaMKII interaction with Shank3 because mutation of the tribasic residue motif in the novel CaMKII-binding domain prevents the rescue of depolarization-induced CREB phosphorylation following knockdown of endogenous Shank3 expression. This leads to the question: why does LTCC-CREB signaling require interactions of CaMKII with both the Cav1.3 NTD AND Shank3? Since a CaMKII holoenzyme can bind simultaneously to multiple CaMKAPs (Robison et al., 2005b), it is possible that different subunits within the same dodecameric CaMKII holoenzyme interact with both the Cav1.3 NTD and Shank3. It is tempting to speculate that such simultaneous interactions could provide a conformational constraint on the cytoplasmic domains of the LTCC, perhaps with functional implications. Alternatively, it is interesting that the initiation of LTCC-CREB signaling was recently shown to require the recruitment of two separate CaMKII holoenzymes to LTCC Ca^2+^ nanodomains and that trans-autophosphorylation between the two holoenzymes is required for the shuttling of CaM to the nucleus (Cohen et al., 2018, Ma et al., 2014). Since trans-autophosphorylation between two separate holoenzymes is highly inefficient in solution (Hanson et al., 1994), another hypothesis is that the two CaMKII-binding sites within the same LTCC complex might bring two holoenzymes in sufficiently close proximity to overcome this biochemical barrier. Indeed, this hypothesis predicts that the loss of any one of these three proteins or disruption of any of their mutual interactions would disrupt the formation of this complex in the LTCC nanodomain, interfering with LTCC-CREB signaling. However, one can readily imagine alternative biochemical mechanisms that might also require multiple CaMKII binding sites within the LTCC nanodomain. Further studies will be needed to precisely define biochemical mechanisms within the LTCC nanodomain that are required to initiate LTCC-CREB signaling.

### CaMKII signaling in Shank3-related neurological disorders

Shank3 is the one of the most commonly mutated genes in individuals diagnosed with autism spectrum disorders (ASD) and is also linked to other neuropsychiatric disorders (Gauthier et al., 2010, Herbert, 2011). Functionally disruptive missense mutations in CaMKIIα and LTCCs have also been identified in patients with ASD and other neurodevelopmental disorders (Akita et al., 2018, Chia et al., 2018, Dick et al., 2016, Kury et al., 2017, Limpitikul et al., 2016, Moon et al., 2018, Nyegaard et al., 2010, Pinggera et al., 2015, Pinggera et al., 2017, Pinggera & Striessnig, 2016, Proietti Onori et al., 2018, Stephenson et al., 2017). In addition, the expression of *c-fos*, a well-studied target of CREB-mediated gene expression, is dysregulated in rodent models of autism (Dubiel & Kulesza, 2015, Orlandini et al., 1996, Williams & Umemori, 2014). However, CREB phosphorylation also may be increased in neuropsychiatric disorders since three unique point mutations in Cav1.3 identified in patients with ASD resulted in gain-of-function phenotypes in various gating mechanisms (Pinggera et al., 2015, Pinggera et al., 2017). Increased CREB phosphorylation has been directly shown with a point mutant in Cav1.2 associated with Timothy Syndrome (Li et al., 2016). Interestingly, increases or decreases in Shank3 expression appear to be associated with distinct neuropsychiatric phenotypes (Bozdagi et al., 2010, Han et al., 2013, Uchino & Waga, 2013). These changes in E-T coupling presumably reflect disruptions in signaling from the LTCC nanodomain to the nucleus that may result from altered interactions between key postsynaptic proteins, such as those linked to ASD (Bourgeron, 2009). Indeed, our recent work showed that CaMKIIα interactions with Shank3 and several other CaMKAPs are disrupted by an ASD-linked *de novo* CaMKIIα E183V mutation (Stephenson et al., 2017), including the Cav1.3 NTD (unpublished observations, Wang & Perfitt). Thus, even though initial studies indicated that genetic disruptions of Shank3 expression failed to detect gross changes in CaMKII (e.g., (Peca et al., 2011)), the present findings suggest that it will be worth testing for more subtle changes in CaMKII signaling following genetic manipulation of Shank3 expression or function, and in animal models of other neuropsychiatric disorders.

## MATERIALS AND METHODS

### Animals

All mice were housed on a 12 h light-dark cycle with food and water ad libitum. CaMKIIα-KO and Camk2a^tm2Sva^ (CaMKIIα^T286A^) mice on a C57B/6J background were as previously described (Giese et al., 1998, Marks et al., 2018). Wild-type (WT) and homozygous littermates were generated using a HetxHet breeding strategy. Both male and female mice age P28-30 were used for biochemical studies. All animal experiments were approved by the Vanderbilt University Institutional Animal Care and Use Committee and were carried out following the US National Institutes of Health Guide for the Care and Use of Laboratory Animals.

### Antibodies used

The following antibodies were used for immunoblotting at the indicated dilutions: mouse anti-CaMKIIα 6G9 (Thermo Fisher Scientific, catalog MA1–048, 1:5000), pT286 CaMKIIα (Santa Cruz Biotechnology, catalog sc-12886-R, 1:3000), mouse anti-Shank3 (University of California at Davis/National Institutes of Health NeuroMab Facility, catalog N367/62, 1:2000), rabbit monoclonal anti-Shank3 (D5K6R) (Cell Signaling, catalog 64555, 1:3000), goat anti-GST (GE Healthcare Life Sciences, catalog 27-4577-01, 1:5000), polyclonal goat CaMKII antibody (RRID: AB_2631234, 1:5000) (Mcneill & Colbran, 1995), mouse anti-PSD-95 (NeuroMab, catalog 75-028, 1:50,000), mouse anti-GFP (Vanderbilt Antibody and Protein Resource catalog 1C9A5, 1:3000), mouse anti-HA (Biolegend, catalog 901503, 1:3000), HRP-conjugated anti-rabbit (Promega, catalog W4011, 1:6000), HRP-conjugated anti-mouse (Promega, catalog W4021, 1:6000), HRP-conjugated anti-goat (Abcam, catalog Ab6741, 1:3000), IR dye-conjugated donkey antimouse 800CW (LI-COR Biosciences, catalog 926–32213, 1:10,000), and IR dye-conjugated donkey anti-goat 680LT (LI-COR Biosciences, catalog 926 – 68022, 1:10,000).

### DNA constructs

A construct encoding GFP-Shank3 was a gift from Dr. Craig Garner (Stanford University). The construct expressing shRNA (pLL3.7) was a gift from Dr. Luk Van Parijs (Massachusetts Institute of Technology). This plasmid was modified to replace the CMV promoter with a 0.4kb fragment of the mouse CaMKII promoter (designated as pLLCK) that is primarily active only in excitatory neurons (Dittgen et al., 2004). Control shRNA (5’-TCGCTTGGGCGAGAGTAAG-3’) was designed following Boudkkazi *et al*. (Boudkkazi et al., 2014). Shank3 shRNA (5’-GGAAGTCACCAGAGGACAAGA-3’) was designed following Verpelli *et al*. (Verpelli et al., 2011). Sequences encoding shRNA were inserted into pLLCK at HpaI and XhoI restriction sites. The Shank3 shRNA-resistant construct (Shank3^R^) was designed by introducing 6 “silent” nucleotide mutations in the target site that do not alter amino acid sequence at Arg^1187^, Lys^1188^, Ser^1189^, and Pro^1190^. The mAp-Shank3 construct was generated by inserting Shank3 cDNA into pmApple-C1 construct at BglII and EcoRI restriction sites. The Shank3 shRNA-resistant construct containing a deletion of the PDZ domain (mAp-Shank3^R-^ΔPDZ) was generated by in-frame PCR deletion of the entire 270bp region encoding ^572^Iso-Val^661^ from mAp-Shank3^R^.

Rat Cav1.3 complete coding sequence (Genbank accession number AF370010) was a gift from Dr. Diane Lipscombe (Brown University). A plasmid encoding Cav1.3-CTD with an N-terminal HA-tag (pCGNH-Cav1.3-CTD, for co-immunoprecipitation) was made by inserting rat Cav1.3 cDNA encoding ^1469^Met-Leu^2164^ into the pCGN vector, a gift from Dr. Winship Herr (Université de Lausanne). All constructs were confirmed by DNA sequencing.

### Mouse forebrain homogenization and subcellular fractionation

Forebrains were dissected and fractionated as previously described (Baucum et al., 2015, Stephenson et al., 2017). Briefly, P28-30 mice were anesthetized with isofluorane, decapitated, and forebrains were quickly dissected, cut in half down the midline, and a half brain was immediately homogenized in an isotonic buffer (150 mM KCl, 50 mM Tris HCl, pH 7.5, 1 mM DTT, 0.2 mM PMSF, 1 mM benzamidine, 1 μM pepstatin, 10 mg/L leupeptin, 1 μM microcystin). The homogenate (~2 ml) was rotated end-over-end at 4°C for 30 min and then centrifuged at 100,000 x *g* for 1 hr. After removing the supernatant (cytosolic S1 fraction), the pellet was resuspended in the isotonic buffer containing 1% (v/v) Triton X-100, triturated until homogeneous, and then rotated end-over-end at 4°C for 30 min. Lysates were then centrifuged at 10,000 x g, and the supernatant (Triton-soluble membrane S2 fraction) was removed. The second pellet (Triton-insoluble synaptic P2 fraction) was resuspended in isotonic buffer containing 1% Triton X-100 and 1% deoxycholate and then sonicated. The P2 fraction was then mixed with 4X SDS-PAGE sample buffer or used for immunoprecipitation studies (see below).

### Recombinant mouse CaMKIIα and GST-tagged protein purification

Expression and purification of recombinant mouse CaMKIIα has been described previously (Mcneill & Colbran, 1995). GST-Shank3 constructs were created by PCR amplification of the relevant cDNA fragments for insertion between EcoR1 and BamH1 restriction sites in pGEX6P-1. GST-GluN2B was described previously (Strack et al., 2000). The vectors encoding GST fusion proteins were transformed into BL21 (DE3) pLysS bacteria cells, and proteins were purified as previously described (Robison et al., 2005a).

### CaMKII autophosphorylation and GST co-sedimentation assays

CaMKIIα (1.25 μM subunit) was incubated on ice for 90 s with 50 mM HEPES, pH 7.5, 10 mM magnesium acetate, 0.5 mM CaCl2, 1 μM CaM, 1 mM DTT, and 400 μM ATP. Autophosphorylation reactions were terminated with 45 mM EDTA. The reaction was then diluted 10-fold using 1X GST pulldown buffer (50 mM Tris-HCl pH 7.5; 200 mM NaCl; 1% (v/v) Triton X-100). Autophosphorylated CaMKIIα (125 nM subunit) was incubated with GST or GST-fusion protein (125 nM) and Pierce Glutathione Agarose beads (Cat. #16101, 10 μL packed resin). Reactions were rocked for 1 hr at 4°C. Beads were washed three times with GST buffer, and proteins were eluted with 20 mM glutathione, pH 8.0 for 10 minutes (Sigma).

### Western blot analysis

Samples were resolved on 10% SDS-PAGE gels and transferred to nitrocellulose membrane (Protran). The membrane was blocked in blotting buffer containing 5% nonfat dry milk, 0.1% Tween 20, in Tris-buffered saline (20 mM Tris, 136 mM NaCl) at pH 7.4 for 30 min at room temperature. The membrane was incubated with primary antibody (see dilutions above) in blotting buffer for 1 hr at room temperature or overnight at 4°C. After washing, membranes were incubated with HRP-conjugated secondary antibody for 30 min at room temperature, washed again, and then visualized using enzyme-linked chemiluminescence using the Western Lightening Plus-ECL, enhanced chemiluminescent substrate (PerkinElmer) and visualized using Premium X-ray Film (Phenix Research Products) exposed to be in the linear response range. Images were quantified using ImageJ software. Secondary antibodies conjugated to infrared dyes (LI-COR Biosciences) were used for development with an Odyssey system (LI-COR Biosciences).

### Cell culture, transfection, and lysis

HEK293T cells (ATCC, catalog CRL-3216) were cultured and maintained in DMEM containing 10% FBS, L-glutamine, and 1% penicillin/streptomycin at 37°C in 5% CO2. Cells plated on 10 cm dishes were transfected with 5-10 μg of DNA. After 24-48 hr, cells were rinsed in ice cold PBS and lysed in ice-cold lysis buffer (150 mM NaCl, 25 mM Tris-HCl, pH 7.5, 1% Triton-X 100, 2 mM EDTA, 2 mM EGTA, 1 mM DTT, 0.2 mM PMSF, 1 mM benzamidine, 10 μg/ml leupeptin, and 10 μM pepstatin).

### Immunoprecipitation

Mouse forebrains fractions or HEK293T cell lysates were incubated with either mouse anti-CaMKIIα (Thermo Fisher Scientific catalog #MA1–048), rabbit anti-Shank3 (Bethyl catalog #A304-178A), or rabbit anti-GFP (Thermo Fisher Scientific, A-11222) and rocked end-over-end at 4°C for 1 hour with 10 μl prewashed Dynabeads Protein A (Thermo Fisher Scientific, Cat. #10001D, for rabbit) or Protein G (Thermo Fisher Scientific, Cat. #10002D, for mouse). HEK293T lysates were supplemented with 1.5 mM CaCl2 and 1 μM calmodulin (final concentrations) to activate CaMKII during immunoprecipitation. The beads were isolated magnetically and washed three times using lysis buffer before eluting proteins using 2X SDS-PAGE sample buffer. Inputs and immune complexes were immunoblotted side by side as indicated (above)

### Colocalization Studies

ST*Hdh*^Q7/Q7^ cells (Trettel et al., 2000), a gift from Dr. Aaron Bowman (Vanderbilt University) were cultured and maintained in DMEM containing 10% FBS, L-glutamine, 1% penicillin/streptomycin, and 400 μg/ml G418 (Mediatech) at 33°C in 5% CO2. Cells were plated onto 15 mm coverslips pre-treated with poly-D-lysine in 12-well plates and transfected with 3 μg of DNA overnight. Media was then removed and cells were incubated in serum-free DMEM containing either 0.49% dimethyl sulfoxide (Pierce) or a differentiation medium (serum-free DMEM supplemented with 10 ng/ml fibroblast growth factor (Promega), 240 μM isobutylmethylxanthine (Sigma), 20 μM 12-0-tetradecanoylphorbol-13-acetate (Sigma), 48.6 μM forskolin (Sigma), and 5 μM DA (Sigma). After 8-14 hours, cells were fixed in ice-cold 4% paraformaldehyde-4% sucrose in 0.1 M Phosphate Buffer pH 7.4 for 3 minutes and −20°C methanol for 10 minutes. Coverslips were mounted on microscope slides using ProLong Gold antifade reagent with DAPI (Thermo Fisher Scientific, catalog P36931). Images were collected using a Zeiss 880 inverted confocal microscope using 63X objective. Thresholding and intensity correlation analysis to compare normalized pixel intensities in each color channel were performed using ImageJ as previous described (Baucum et al., 2010). GFP and mApple channels were automatically thresholded before calculating the intensity correlation quotient (ICQ). This method interprets ICQ values in the following ranges: 0 < ICQ ≤ +0.5, dependent overlap of fluorescent signals; ICQ = 0, random overlap of fluorescent signals; and 0 > ICQ ≥ −0.5, segregation of fluorescent signals (Li et al., 2004).

### Primary hippocampal neuronal cultures and neuronal stimulation assay

Dissociated hippocampal neurons were prepared from E18 Sprague Dawley rat embryos, as previously described (Shanks et al., 2010). Neurons were transfected at 7-9 days *in vitro* (DIV) using Lipofectamine 2000 following the manufacturer’s directions (Thermo Fisher Scientific). A total of 1 μg of DNA was transfected for each well of a 12-well plate for 2-3 hours before switching back to conditioned media. At DIV 13-14, neurons were pre-incubated for 2hr with 5K Tyrode’s solution (150 mM NaCl, 5 mM KCl, 2 mM CaCl_2_, 2 mM MgCl_2_, 10 mM glucose and 10 mM HEPES pH 7.5 (~313 mOsm)) with 1 μM TTX, 10 μM APV and 50 μM CNQX to suppress intrinsic neuronal activity by blocking sodium channels, NMDA receptors and AMPA receptors, respectively. Neurons were then treated with either 5K Tyrode’s or 40K Tyrode’s solution (adjusted to 40 mM KCl and 115 mM NaCl, with all three inhibitors present) for 90 seconds and fixed using ice-cold 4% paraformaldehyde-4% sucrose in 0.1 M Phosphate Buffer pH 7.4 for 3 minutes and −20°C methanol for 10 minutes. Neurons were washed three times with PBS, permeabilized with PBS+0.2% Triton X-100, and then incubated with blocking solution for one hour (1X PBS, 0.1% Triton X-100 (v/v), 2.5% BSA (w/v), 5% Normal Donkey Serum (w/v), 1% glycerol (v/v)). Neurons were then incubated with blocking solution overnight with primary antibodies: rabbit anti-pCREB (Cell Signaling, catalog #9198, 1:1000), and mouse anti-CaMKIIα 6G9 (Thermo Fisher Scientific, catalog MA1–048, 1:1000). The following day, neurons were washed three times in PBS+0.2% Triton X-100, then incubated with blocking solution for 1 hour with secondary antibodies: donkey anti-rabbit 647 Alexa Fluor 647 (catalog A-31573) and donkey anti-mouse Alexa Fluor 546 (catalog A-10036). Neurons were washed with PBS three times and mounted on slides using Prolong Gold Antifade Mountant with DAPI (Thermo Fisher Scientific, catalog P36931).

### Neuronal imaging and quantification

The experimenter was blinded to the transfection conditions by coding the culture dishes prior to microscopy and image analysis. Images were collected using a Zeiss 880 inverted confocal microscope with a 40x/1.30 Plan-Neofluar oil lens. The binocular lens was used to identify transfected neurons based on EGFP expression from the shRNA construct. In experiments with mAp-Shank3^R^ rescue, mAp expression was also confirmed in EGFP-positive cells. The DAPI channel was then used to focus on the z-plane that yielded the highest DAPI signal (one that presumably runs through the nuclei) for image acquisition. Images were then collected in all channels and MetaMorph Microscope Automation and Image Analysis Software (Molecular Devices) was used to quantify pCREB or c-fos signals. Briefly, nuclei were identified by thresholding the DAPI channel to create and select the nuclear regions of interest (ROIs). The ROIs were then transferred to other channels to measure the average pCREB or c-fos intensity. ROIs were collected from transfected (EGFP-positive) neurons and nearby non-transfected (EGFP-negative) neurons. The relative intensity was calculated as [(channel^x^-channel^5K^)/(channel^40K^-channel^5K^)], where channel^x^ is the signal being calculated, and channel^5K^ and channel^40K^ are the average signals of the 5K and 40K conditions in that batch of cultured neurons, respectively. Data shown were collected from images of the indicated total number of neurons from 3-5 independent cultures.

## ACKNOWLEDGEMENTS

We thank Drs. Craig Garner, Luk Van Parijs, Diane Lipscombe, and Winship Herr for generously providing various plasmids, as detailed above. Confocal imaging and analysis were performed in part through the use of the Vanderbilt Cell Imaging Shared Resource (supported by National Institutes of Health Grants CA68485, DK20593, DK58404, DK59637, and EY08126).

This work was supported in part by National Institutes of Health Grants R01-MH063232 and R01-NS078291 (to R. J. C.), and R01-HD061543 (to T. N.), and American Heart Association predoctoral fellowships 18PRE33960034 (to T. L. P.) and 14PRE18420020 (to X. W.). The content is solely the responsibility of the authors and does not necessarily represent the official views of the National Institutes of Health.

## AUTHOR CONTRIBUTIONS

T.L.P. helped conceive the studies, designed experiments, performed most of the experiments, and wrote drafts of the manuscript. T.L.P., X.W., J.R.S. and T.N generated reagents. T.N. and X.W. provided advice concerning experimental design and data analysis/interpretation. R. J. C. conceived and coordinated the studies, and helped to design experiments, analyze/interpret the data, and write the manuscript. All authors reviewed the results, helped edit the manuscript, and approved the final version.

## CONFLICT OF INTEREST

The authors declare no competing financial interests.

---

**Expanded View Figure 1.**
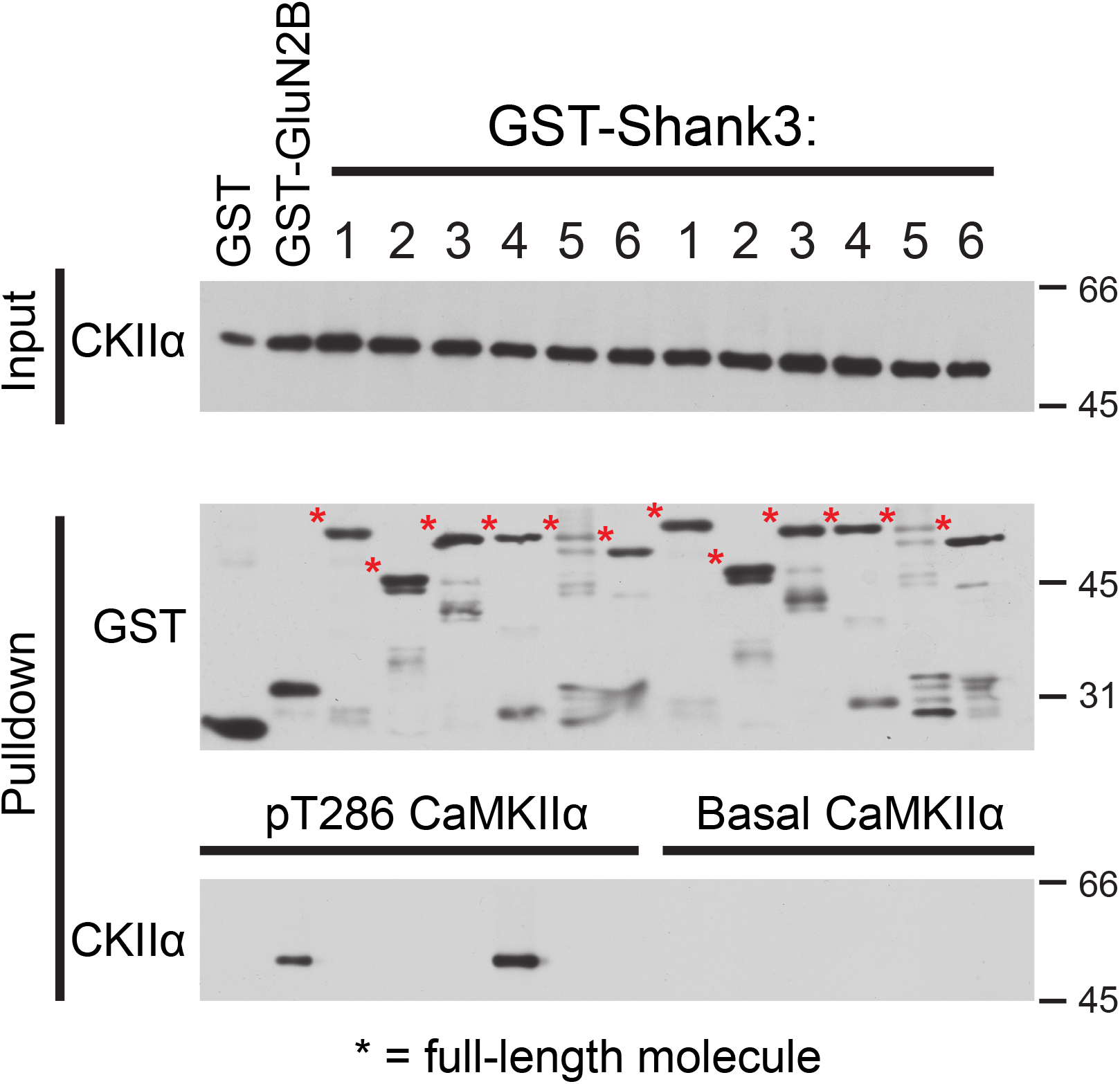
Inactive CaMKIIα does not bind to Shank3. Glutathione agarose co-sedimentation assay with pre-activated (Thr286-phosphorylated) CaMKIIα (left) and basal, inactive CaMKIIα (right). Pre-activated CaMKIIα specifically binds to GST-Shank3 #4 (829-1130) and positive control GST-GluN2B (1260-1309), but not any other fusion proteins. Inactive CaMKIIα does not bind to any GST-Shank3 fusion proteins. Full-length GST fusion proteins denoted with asterisks on the GST immunoblot. Immunoblots are representative of 3 biological replicates.

**Expanded View Figure 2.**
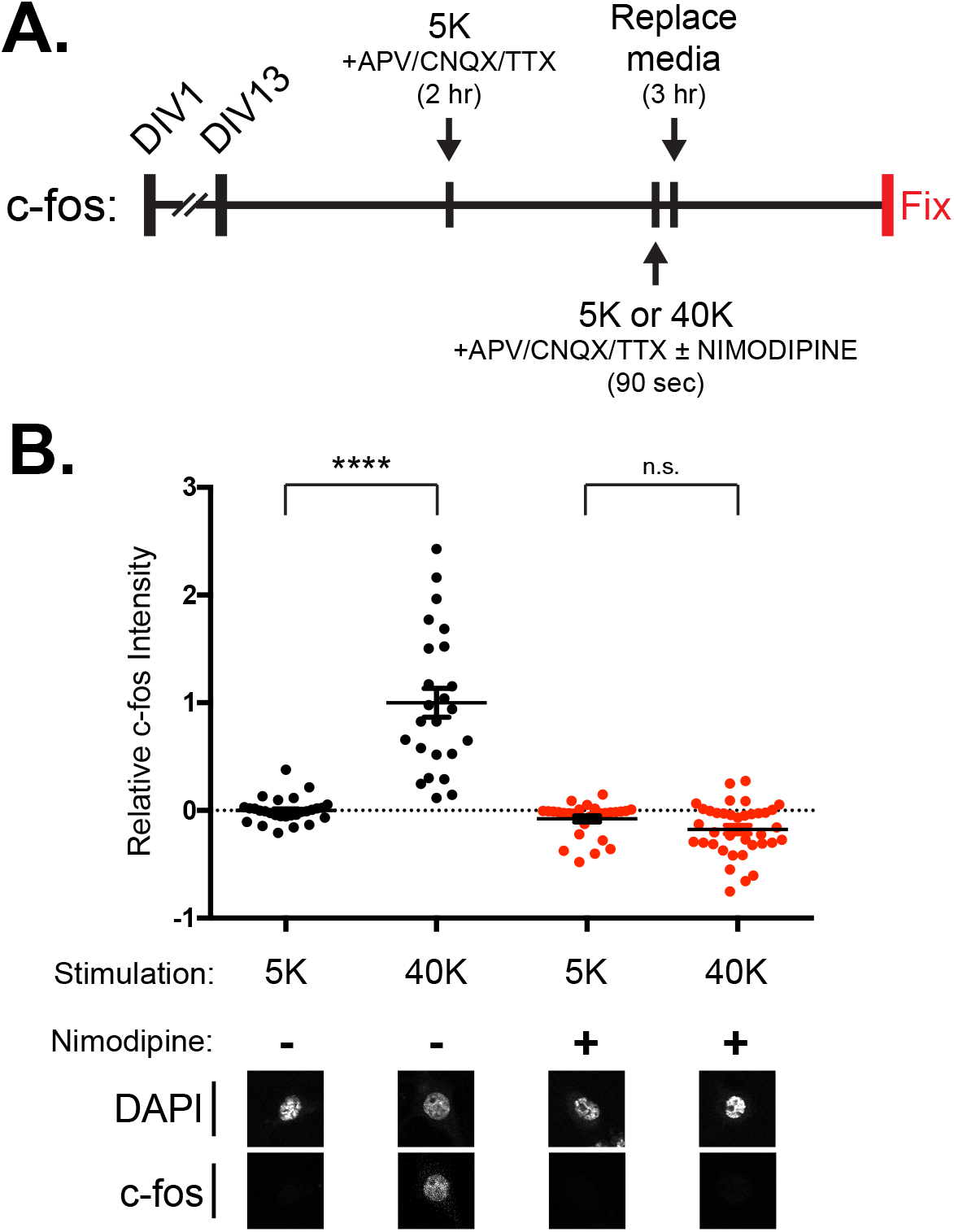
LTCC-CREB signaling increases c-fos expression in neurons. **A.** Schematic of experimental protocol. Primary hippocampal neurons (DIV13) were incubated with inhibitors of NmDa- and AMPA-type glutamate receptors and voltage-gated sodium channels (APV, CNQX, and TTX) in 5 mM KCl (5K) Tyrode’s solution for 2 hours, and then treated with 5K (control) or 40 mM KCl (40K: depolarizing) Tyrode’s solution for 90 seconds before being replaced with original media for 3 hours before fixation. Separate dishes were treated in parallel with selective LTCC blocker nimodipine during stimulation. Neurons were then stained using antibodies to c-fos and DAPI. **B.** The 40K stimulation induced a robust increase in nuclear c-fos expression, which is significantly reduced in neurons treated with nimodipine (2-way ANOVA with two factors (Stimulation, Nimodipine treatment), Stimulation F_(1,114)_ = 51.00, Nipodipine treatment F_(1,114)_ = 98.87, Interaction F_(1,114)_ = 75.87, all p < 0.0001. Tukey’s post-hoc test, **** p < 0.0001). Each data point represents analysis of a single cell accumulated from 2 independent neuronal cultures. All confocal images show a 40 x 40 μm area. The bar graph reports the mean ± SEM.

**Expanded View Figure 3.**
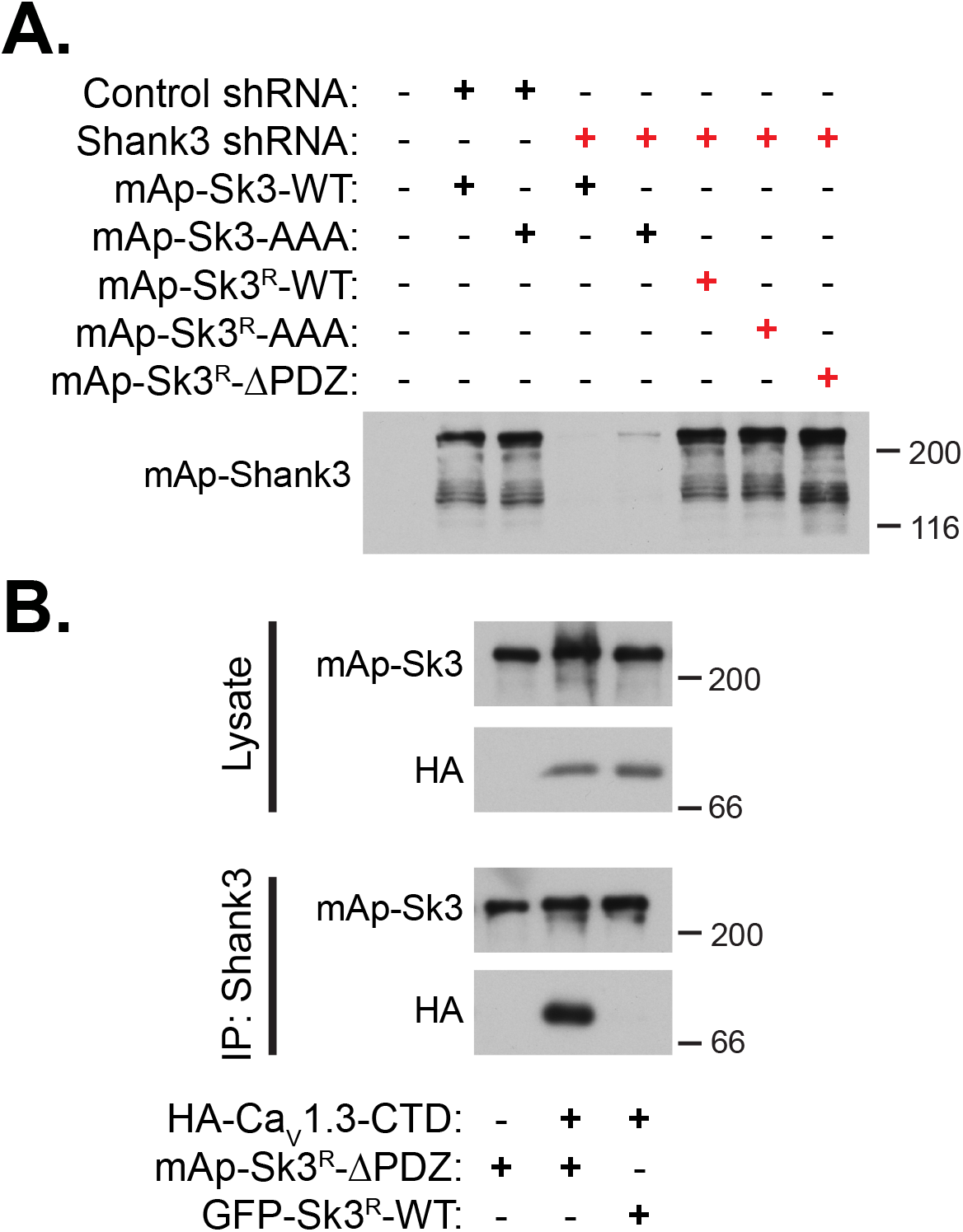
Validation of mAp-Shank3^R^ shRNA-resistance and loss of binding to Cav1.3 in mAp-Shank3^R^-ΔPDZ. **A.** Expression of shRNA and mAp-Shank3 constructs in HEK293T cells. Cells were co-transfected with control shRNA or shRNA against Shank3, along with mAp-Shank3 constructs that are not resistant to the shRNA (mAp-Shank3-WT and mAp-Shank3-AAA) or contain ‘silent’ mutations that imbue shRNA resistance (mAp-Shank3^R^ constructs). Immunoblotted with NeuroMab Shank3 antibody. B. Soluble fractions of HEK293T cells expressing HA-Cav1.3-CTD with mAp-Shank3^R^-WT or mAp-Shank3^R^-ΔPDZ were immunoprecipitated using a Shank3 antibody. Co-precipitation of HA-Cav1.3-CTD is observable with mAp-Shank3^R^-WT, but not with mAp-Shank3^R^-ΔPDZ. Immunoblots are representative of 2 biological replicates.

